# CGAN-Cmap: protein contact map prediction using deep generative adversarial neural networks

**DOI:** 10.1101/2022.07.26.501607

**Authors:** Mohammad Madani, Mohammad Mahdi Behzadi, Dongjin Song, Horea Ilies, Anna Tarakanova

## Abstract

Protein contact maps represent spatial pairwise inter-residue interactions, providing a protein’s translationally and rotationally invariant topological representation. Accurate contact map prediction has been a critical driving force for improving protein structure prediction, one of computational biology’s most challenging problems in the last half-century. While many computational tools have been developed to this end, most fail to predict accurate contact maps for proteins with insufficient homologous protein sequences, and exhibit low accuracy for long-range contacts. To address these limitations, we develop a novel hybrid model, CGAN-Cmap, that uses a generative adversarial neural network embedded with a series of modified squeeze and excitation residual networks. To exploit features of different dimensions, we build the generator of CGAN-Cmap via two parallel modules: sequential and pairwise modules to capture and interpret distance profiles from 1D sequential and 2D pairwise feature maps, respectively, and combine them during the training process to generate the contact map. This novel architecture helps to improve the contact map prediction by surpassing redundant features and encouraging more meaningful ones from 1D and 2D inputs simultaneously. We also introduce a new custom dynamic binary cross-entropy (BCE) as the loss function to extract essential details from feature maps, and thereby address the input imbalance problem for highly sparse long-range contacts in proteins with insufficient numbers of homologous sequences. We evaluate the performance of CGAN-Cmap on the 11th, 12th, 13th, and 14th Critical Assessment of protein Structure Prediction (CASP 11, 12, 13, and 14) and CAMEO test sets. CGAN-Cmap significantly outperforms state-of-the-art models, and in particular, it improves the precision of medium and long-range contact by at least 3.5%. Furthermore, our model has a low dependency on the number of homologous sequences obtained via multiple sequence alignment, suggesting that it can predict protein contact maps with good accuracy for those proteins that lack homologous templates. These results demonstrate an efficient approach for fast and highly accurate contact map prediction toward construction of protein 3D structure from protein sequence.

**Data availability:** All datasets and source codes are provided in: https://github.com/mahan-fcb/CGAN-Cmap-A-protein-contact-map-predictor

## Introduction

Protein structure determination from sequence has long been one of the most challenging problems in structural biology^1,2,3^. Experimental approaches to determine the 3D structure of a protein are often time-consuming and costly^4,5^. Only about 200,000 protein structures have been identified through experimental methods such as X-ray crystallography compared to millions of existing protein sequences^6^. To bridge the sequence-structure gap, ab initio computational methods play pivotal complementary roles in resolving protein structure and function^2,7^. Progress in community-wide Critical Assessment of Structure Prediction (CASP) experiments has demonstrated that contact map prediction, used as an intermediary constraint, boosts the accuracy of ab initio folding simulations and improves the success rate for prediction of protein targets^8^. Knowledge about contacts between pairs of residues provides a translationally and rotationally invariant topological representation of a protein that can capture the global topology of the protein fold^8,9^.

A number of computational tools to predict protein contact maps have been developed, most falling under two common classes of methods: Evolutionary Coupling Analysis (ECA) and machine learning (ML)^10^ methods. ECA predicts contact maps by analyzing evolutionary correlations of proteins derived from multiple sequence alignments (MSAs). Direct evolutionary analysis (DCA) is most accurate among ECA methods, using MSAs to identify direct evolutionary residue-residue relationships and correlations. DCA considers the effect of residues in different positions on each pair of residues to quantify the correlations between them^11,12^. Popular DCA methods include CCMPred^13^, FreeContact^14^, GREMLIN^15^, and PSICOV^16^. To find these relationships, DCA commonly utilizes graphical lasso^11^ and pseudo-likelihood maximization^12^. Graphical lasso finds the graph structure from a covariance matrix, and pseudo-likelihood maximization is a probabilistic model to find the strength of interactions between residues^11,12^. Notably, DCA methods generally exhibit two major limitations. First, DCA-based methods have poor performance when the number of homologous sequences is lower than approximately 50^17,18^. Second, these methods extract only linear relationships between pairs of residues^17^. Conversely, the relationships between pairs of residues are intrinsically non-linear^17^.

By contrast, machine learning, and more specifically deep learning approaches, have been successfully utilized to find more accurate contact maps for proteins with few homologous sequences^19^. Most of the recently proposed models tested in CASP are based on deep learning, in particular residual neural networks (ResNets)^17,18,20^, which resolve the classical machine learning vanishing gradient problem, and more importantly, show sufficient depth to accurately predict protein contact maps^21^. ResNets effectively encourage most important features and bypass low quality information within feature maps. Notably, two of the most commonly used and accurate contact map predictors (as confirmed by CASP 12 and 13), RaptorX-contact^20^ and TripletRes^17^, are ResNet-based. Besides residual networks, other deep learning models show comparable performance in predicting accurate contact maps, especially for long-range contacts^22,23^. Generative Adversarial Networks (GANs) are one of a powerful class of neural networks used for unsupervised learning, capable of learning low and high-level patterns^24^. GANs comprise two competitive networks, namely a generator network and a discriminator. During training, the generator learns to maximize the accuracy of generated samples to fool the discriminator. On the other hand, the discriminator tries to distinguish between the generated samples and real data^25^. Conditional GANs have been widely adopted for high-level resolution generation tasks including image-to-image translation^26^. Thus, this functionality enables GANs to be used in making predictions of optimal and more accurate contact maps^24,27,28^.

Despite recent progress in contact map prediction, a challenging outstanding problem remains: the prediction of medium and long-range contacts (i.e., where the distance between position indices of pairs of amino acid residues is more than 12 Å). These types of contacts are highly sparse, complicating accurate prediction^17^. Sparsity creates an imbalance problem, when the ratio of contacting and uncontacting residues is low (less than 3%). To tackle this, we build a new tool to accurately predict contact maps for medium and long-range contacts regardless of the number of homologous sequences available. Inspired by the capabilities of GANs for generating high-resolution images from input features, in this study we propose a contact map predictor, CGAN-Cmap, constructed via integration of a modified squeeze excitation residual neural network (SE-ResNet), SE-Concat, and a conditional GAN. In the architecture of the GAN generator, we define two novel subnets to extract feature representations during training and feed them to subsequent layers gradually. We also introduce a custom loss function for training the model, a dynamic weighted binary cross-entropy (BCE) loss function, which assigns a dynamic weight for classes based on the ratio of the uncontacted class to the contacted class in each iteration of the training process. This loss function emphasizes misclassified residue pairs to enhance the model training and tackle the imbalance problem. By taking advantage of the model architecture and custom loss function, the receptive field and feature reusability are significantly enhanced, which allows us to capture more complex non-linear relationships between amino acid residues even for proteins with few homologous sequences. We assess the performance of CGAN-Cmap on existing recent CASP datasets (CASP 11, CASP 12, CASP 13, CASP 14). The results reveal that CGAN-Cmap utilizing the dynamic BCE as a custom loss function consistently outperforms leading existing models^17,20^ on medium and long-range contacts for all recent CASP datasets. We also show that CGAN-Cmap has a moderate dependency on the number of homologous sequences available in MSAs, ensuring that it can accurately predict contact maps for proteins with insufficient homologous sequences. Overall, we show that CGAN-Cmap derives its superior performance from a custom loss function to overcome the sparsity problem, and a novel architecture, which increases the receptive field and feature reusability as well as encourages more important features within the feature map.

## Materials and Methods

### Contact map definition

The protein contact map is symmetric and represents the probability of contact between two residues. If the Euclidean distance is less than 8 Å between two beta carbon (Cβ) atoms within two residues (for glycine alpha carbons (Cα) are used), these residues are considered to be contacting residues in the protein contact map. Generally, contacts are divided into three categories based on the difference, *Diff*, in the position index of two residues, A and B, in the protein sequence, as follows:

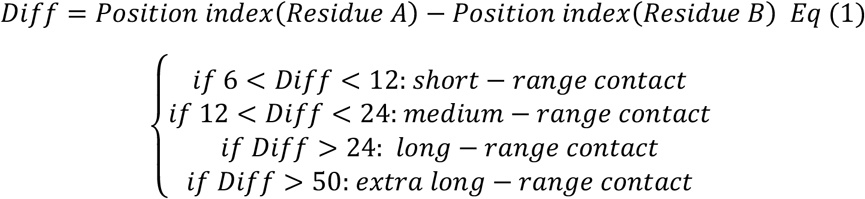

Note that because of the importance of long-range contacts, we introduce a new category termed long-range contacts.

### Datasets

The training dataset for CGAN-Cmap is collected from SCOPe-2.07^29^ and a subset of the Protein Data Bank filtered at 25% sequence identity (PDB25)^30^. The protein sequence selection is conducted per the following criteria: 1) sequence length is between 25-700 residues; 2) the maximum sequence identity is set to 30% to remove redundant sequences via CD-HIT^31^; 3) the native contact map resolution is more than 2.5 Å; 4) the protein contains a single chain. Based on these criteria, 7,128 protein sequences are selected as the training dataset. To evaluate the performance of our model, we use five different blind public test sets including CAMEO (76 proteins)^32^, CASP 11 (110 protein targets)^33^, CASP 12 (54 protein targets)^33^, CASP 13 (41 protein targets)^33^, and CASP 14 (40 protein targets)^33^. In a second redundancy removal process, we eliminate sequences in our initial training set which have sequence identity with our independent test sets of more than 25% via CD-HIT^31^. Supplementary Table S1 summarizes our final training set and independent test sets. The final training set is divided into five subsets. We randomly select one subset as the validation set and leave the remaining subsets as the training set for hyper-parameter tuning. After hyper-parameter tuning, the final model is the average of five models, where each of the five models is trained as described.

### Feature extraction

To build a machine learning model, we extract both pairwise (co-evolution) and sequential features in a process similar to that introduced in RaptorX-Contact^20^. Pairwise features are captured from multiple sequence alignments (MSAs). To generate the MSA for training and test sets, we use the modified version of DeepMSA first proposed by Zhang et al^34^. Additional details on MSA generation are described in the Supplementary Information (SI). After MSA generation, pairwise features including the evolutionary coupling matrix generated by CCMpred^13^, the mutual information matrix^35^, the average product correction (APC)-corrected mutual information matrix^36^, and the pairwise contact potentials matrix^37^ are extracted from MSAs. For sequential features, we use 3- and 8-state protein secondary structure (SS3, SS8), 3-state solvent accessibility (ACC), and features derived from MSAs in a 20-dimensional position-specific scoring matrix (PSSM) and 20-dimensional position-specific frequency matrix (PSFM)^38^. Because PSFM contains the frequency of amino acids in the protein sequence, we include it in the input features to complement the position-specific scoring matrix. In total, sequential features are represented in a two-dimensional matrix with a size of *L* × 54, and pairwise features are represented in a three-dimensional matrix with a size of *L* × *L* × 5, where *L* is the sequence length.

### CGAN-Cmap architecture

We employ a conditional generative adversarial neural network (CGAN) which consists of a generator to produce the contact map and a discriminator to compare the generated contact map with the real contact map (Figure 1). The overall structure of our GAN is inspired by the GANTL model^39^. The generator contains three novel subnets. The first subnet, the conversion subnet, includes a conversion block that converts 1D features to 2D matrices. Unlike the outer concatenation approach used in most existing models^18,20^ to convert 1D features to 2D matrices, our conversion method eliminates the repetition of information, and more importantly, it is learned during the training process, rather than preprocessed. To do so, the model learns the best-converted matrix for each 1D feature (SI Figure S2). The second subnet is the synthesis subnet and includes synthesis (SI Figure S3A) and upsampling (SI Figure S3C) blocks, as explained in the SI. This subnet is responsible for generating the contact map of proteins from the converted 2D matrix from the first subnet and a noise vector. The third subnet is the feature extraction subnet, which receives the 2D features (pairwise features) as input and extracts useful feature maps from 2D information. This subnet includes a series of SE-Concat blocks (SI Figure S3B), that pass the extracted feature map to the synthesis subnet gradually to generate the protein contact map^40,41^. SE-Concat is inspired from the SE-ResNet architecture, modified to use concatenation instead of summation, to reuse the extracted features and increase the information flow between layers^40,42,43^. The squeeze excitation (SE) block (SI Figure S3D) within the SE-Concat block emphasizes the important information and improves channel interdependencies^41^. Unlike other models which treat the feature maps equally, the SE block weighs information based on its effect on the final prediction, to boost model performance.

**Figure 1.**
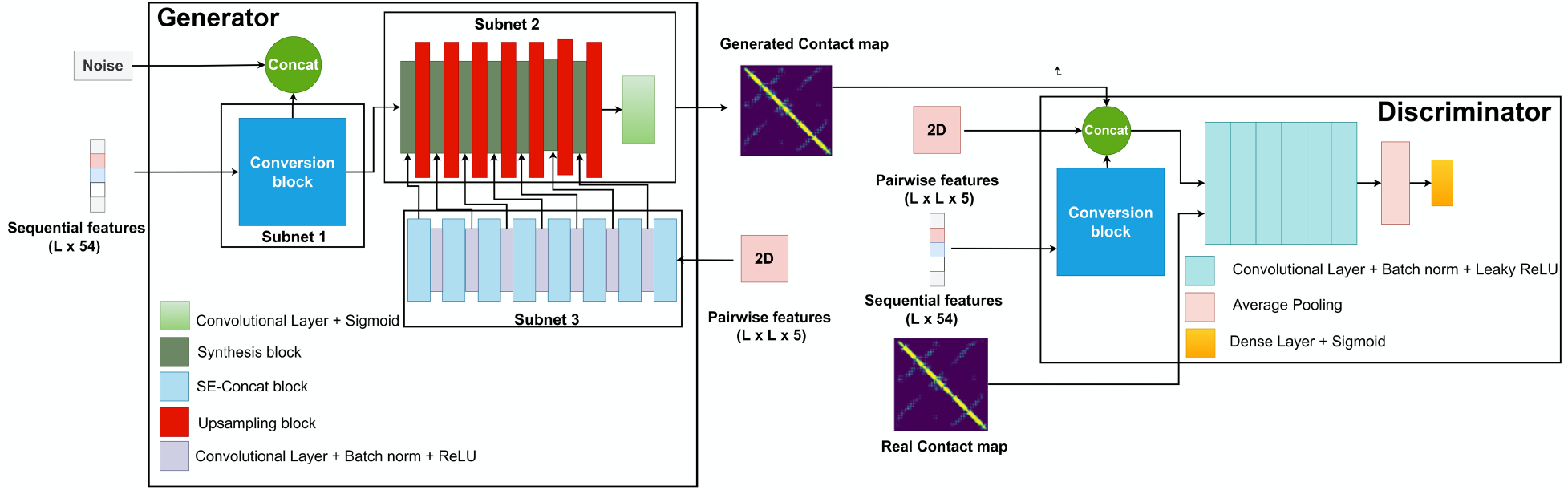
CGAN-Cmap model including generator and discriminator architecture.

The discriminator consists of four convolutional layers and a dense layer that determine the probability of the image being real. The input to the discriminator is the original 2D features, converted 2D features derived from 1D features, and their corresponding contact map. It is worth noting that the discriminator uses the output of the dot layer in the conversion block as the converted 2D features, and ignores the remaining three convolutional layers in the conversion block. This ensures that all matrices have the same dimension, and the output of the dot layer has the same size as the contact map, so we can concatenate the converted 2D features and the original 2D features with the contact map.

### Training process

We aim to train an end-to-end model in a single training process. To do this, we use mini-batches, because the training samples have different size. We choose a batch of training samples and then consider the largest sample size in the batch; then, we pad the samples using the zero-padding method. The padding size is determined based on the number of down samplings in the feature extraction subnet. Our GAN is trained in three steps. In the first step, we train the discriminator on the real data. Then, we repeat the same process for the fake samples generated by the generator. In the final step, we freeze the discriminator and train the generator on the training data set. The hyper-parameters used in the training process are selected based on the grid search. We use the ADAM optimizer and the binary cross-entropy (BCE) loss function for the discriminator.

Selecting a suitable loss function for the generator is essential, since training a deep learning model for contact prediction is the process to determine the parameters of the network by minimizing the loss between the predicted protein contact maps and the real protein contact maps in the training set. Protein contact map prediction is highly imbalanced, as the ratio of the number of ones (contacting residues) to the number of zeros (uncontacting residues) is less than 0.3. To identify the appropriate loss function to overcome this imbalance problem, we train the generator using two different loss functions and compare their performance. The first loss function is a standard BCE (*L_bce_*) and the second one is a dynamic weighted BCE (L_wbce_). The loss functions are defined as follows:

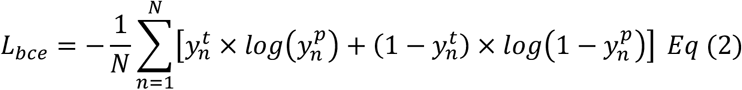

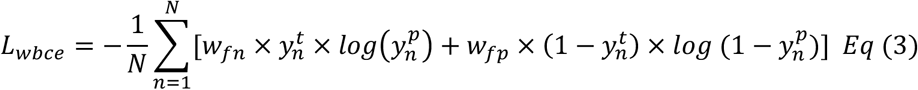

where N is the number of samples, y_n_^t^ is the nth ground truth sample, y_n_^p^ is the n_th_ predicted sample, w_fn_ is the marginal cost of a false negative over true positive, and w_fp_ is the marginal cost of a false positive over true negative^44^. The weights control the contribution of each part of the loss function in the training process. For the dynamic weighted binary cross-entropy loss function, we allow the model to choose the weights of each sample dynamically based on the false negative and false positive values in that sample. Finally, we use five-fold cross-validation for training and the overall performance is calculated by averaging from all folds. Hyperparameters are reported in SI Table S2.

### Model evaluation

We compare the performance of our model to four state-of-the-art contact map predictors, including TripletRes^17^, RaptorX-contact^20^, DeepCon^45^, and DeepCov^46^. We use mean precision of top L/k contacts (k = 10, 5, 2, and 1; L is the sequence length) for short, medium, long, and extra long-range contacts to evaluate the prediction performance of the protein contact map tools.

## Results & Discussion

### Model performance on CASP 11, 12, and CAMEO targets

Table 1 summarizes the performance of our proposed tool on CASP 11, CASP 12, and CAMEO test sets, with standard BCE and dynamic weighted BCE, compared with five existing models. The results show that our models with both standard BCE and dynamic weighted BCE outperform state-of-the art predictors when performance is evaluated based on mean precision. For instance, on CASP 12, mean precision of CGAN-Cmap for long-range top L/10, L/5, and L/2 contacts is higher by at least 4%, 5%, and 4%, respectively, compared to close competitors. These results confirm that the combination of the GAN model with our modified version of residual network blocks (SE-Concat blocks) and dynamic BCE loss function can be employed to improve model performance for protein contact map prediction. For the CAMEO dataset, CGAN-Cmap shows performance slightly inferior (by less than 1%) than TripletRes for some contact ranges. We expect this result to reflect the nature of proteins within the CAMEO dataset: membrane proteins found in the CAMEO dataset are more difficult prediction targets compared to globular proteins.

**Table 1.** Model performance of CGAN-Cmap, TripletRes^17^, RaptorX-contact^20^, DeepCon^45^, and DeepCov^46^ based on the mean precision of short, medium, and long-range top L/10, L/5, L/2, and L contacts in predicted contact maps for CASP 11^33^, CASP 12^33^, and CAMEO^32^ test sets. The highest accuracy in each category is highlighted in bold font.

| Test set | Model | Short-range |  |  |  | Medium-range |  |  |  | Long-range |  |  |  |
| --- | --- | --- | --- | --- | --- | --- | --- | --- | --- | --- | --- | --- | --- |
|  |  | L | L/2 | L/5 | L/10 | L | L/2 | L/5 | L/10 | L | L/2 | L/5 | L/10 |
| CASP 11 | DeepCon | 0.21 | 0.40 | 0.70 | 0.77 | 0.33 | 0.50 | 0.71 | 0.81 | 0.47 | 0.64 | 0.72 | 0.72 |
|  | DeepCov | 0.21 | 0.39 | 0.69 | 0.80 | 0.32 | 0.50 | 0.71 | 0.79 | 0.49 | 0.62 | 0.72 | 0.77 |
|  | RaptorX-contact | 0.27 | 0.46 | 0.74 | 0.83 | 0.35 | 0.56 | 0.77 | 0.86 | 0.56 | 0.69 | 0.77 | 0.82 |
|  | TripletRes | 0.26 | 0.44 | 0.72 | 0.83 | 0.35 | 0.57 | 0.75 | 0.85 | 0.57 | 0.68 | 0.75 | 0.83 |
|  | CGAN-Cmap (standard) | 0.25 | 0.42 | 0.72 | 0.81 | 0.36 | 0.52 | 0.75 | 0.83 | 0.55 | 0.69 | 0.74 | 0.82 |
|  | CGAN-Cmap (dynamic) | **0.27** | **0.49** | **0.74** | **0.87** | **0.38** | **0.57** | **0.79** | **0.88** | **0.60** | **0.72** | **0.83** | **0.85** |
| CASP 12 | DeepCon | 0.25 | 0.58 | 0.69 | 0.74 | 0.27 | 0.59 | 0.62 | 0.72 | 0.35 | 0.55 | 0.60 | 0.64 |
|  | DeepCov | 0.23 | 0.62 | 0.70 | 0.76 | 0.33 | 0.64 | 0.66 | 0.77 | 0.33 | 0.59 | 0.63 | 0.68 |
|  | RaptorX-contact | 0.44 | 0.69 | 0.75 | 0.82 | 0.48 | 0.69 | 0.72 | 0.82 | 0.47 | 0.69 | 0.77 | 0.79 |
|  | TripletRes | 0.47 | 0.70 | 0.77 | 0.80 | 0.46 | 0.72 | 0.76 | 0.83 | 0.45 | 0.67 | 0.71 | 0.77 |
|  | CGAN-Cmap (standard) | 0.53 | 0.68 | 0.80 | 0.86 | 0.49 | 0.77 | 0.79 | 0.86 | 0.55 | 0.70 | 0.80 | 0.82 |
|  | CGAN-Cmap (dynamic) | **0.58** | **0.72** | **0.81** | **0.87** | **0.50** | **0.67** | **0.80** | **0.86** | **0.59** | **0.73** | **0.81** | **0.83** |
| CAMEO | DeepCon | 0.18 | 0.33 | 0.48 | 0.55 | 0.22 | 0.34 | 0.56 | 0.58 | 0.34 | 0.46 | 0.53 | 0.57 |
|  | DeepCov | 0.19 | 0.31 | 0.49 | 0.57 | 0.19 | 0.34 | 0.53 | 0.59 | 0.32 | 0.48 | 0.51 | 0.58 |
|  | RaptorX-contact | 0.24 | 0.39 | 0.58 | 0.67 | 0.29 | 0.43 | 0.63 | 0.70 | 0.42 | 0.55 | 0.65 | 0.69 |
|  | TripletRes | 0.26 | **0.42** | 0.60 | 0.70 | **0.34** | **0.46** | 0.67 | **0.75** | 0.46 | **0.59** | 0.69 | **0.76** |
|  | CGAN-Cmap (standard) | 0.21 | 0.37 | 0.58 | 0.66 | 0.29 | 0.42 | 0.62 | 0.66 | 0.43 | 0.54 | 0.66 | 0.72 |
|  | CGAN-Cmap (dynamic) | **0.27** | 0.41 | **0.61** | **0.72** | 0.33 | 0.45 | **0.69** | 0.74 | **0.47** | 0.57 | **0.70** | 0.75 |

### Model performance on CASP 13 and CASP 14 targets

We also evaluate the performance of our model on recent CASP targets, CASP 13 and CASP 14, and compare to existing models (Table 2). As a note, we extract input features for all models from the same MSA for a fair comparison. In CASP 13, a new range of contact, termed extra long-range, was introduced. So, to be consistent with the recent evaluation protocol in CASP, we evaluate the performance of all models by calculating the mean precision of top L, L/2, L/5, and L/10 contacts in the (medium + long)-range, long-range, and extra long-range. We find that both versions of the CGAN-Cmap model consistently outperform all existing models for all top L/k contacts for k=1, 2, 5, 10, in the (medium + long)-range, long-range, and extra long-range. For example, the mean precision of CGAN-Cmap for top L contacts is 1.4% and 0.9% higher than TripletRes and RaptorX-contact, respectively. For long and (medium + long)-range contacts, the gap between the mean precision of CGAN-Cmap and both RaptorX-contact and TripletRes is also substantial (between 2% - 5.5%). For extra-long range contacts prediction, CGAN-Cmap model achieves mean precision at least 3% higher than close competitors. However, in CASP 13, for top L/10 and L/5 extra long-range contacts, the performance of CGAN-Cmap is slightly inferior to TripletRes and RaptorX-contact (by 1.2% for L/5 and 1.9% for L/10). As a case study, we also compare the prediction performance of CGAN-Cmap on all individual targets within CASP 12, CASP 13, and CASP 14 with TripletRes^17^ and RaptorX-contact^20^(Figure 2). CGAN-Cmap has higher precision values across most of the targets, for top L/2 and L/5 (medium + long)-range, long-range, and extra long-range contacts in comparison to state-of-the-art models, despite CGAN-Cmap’s reduced training set size, number of layers in the architecture, and trainable parameters.

**Figure 2.**
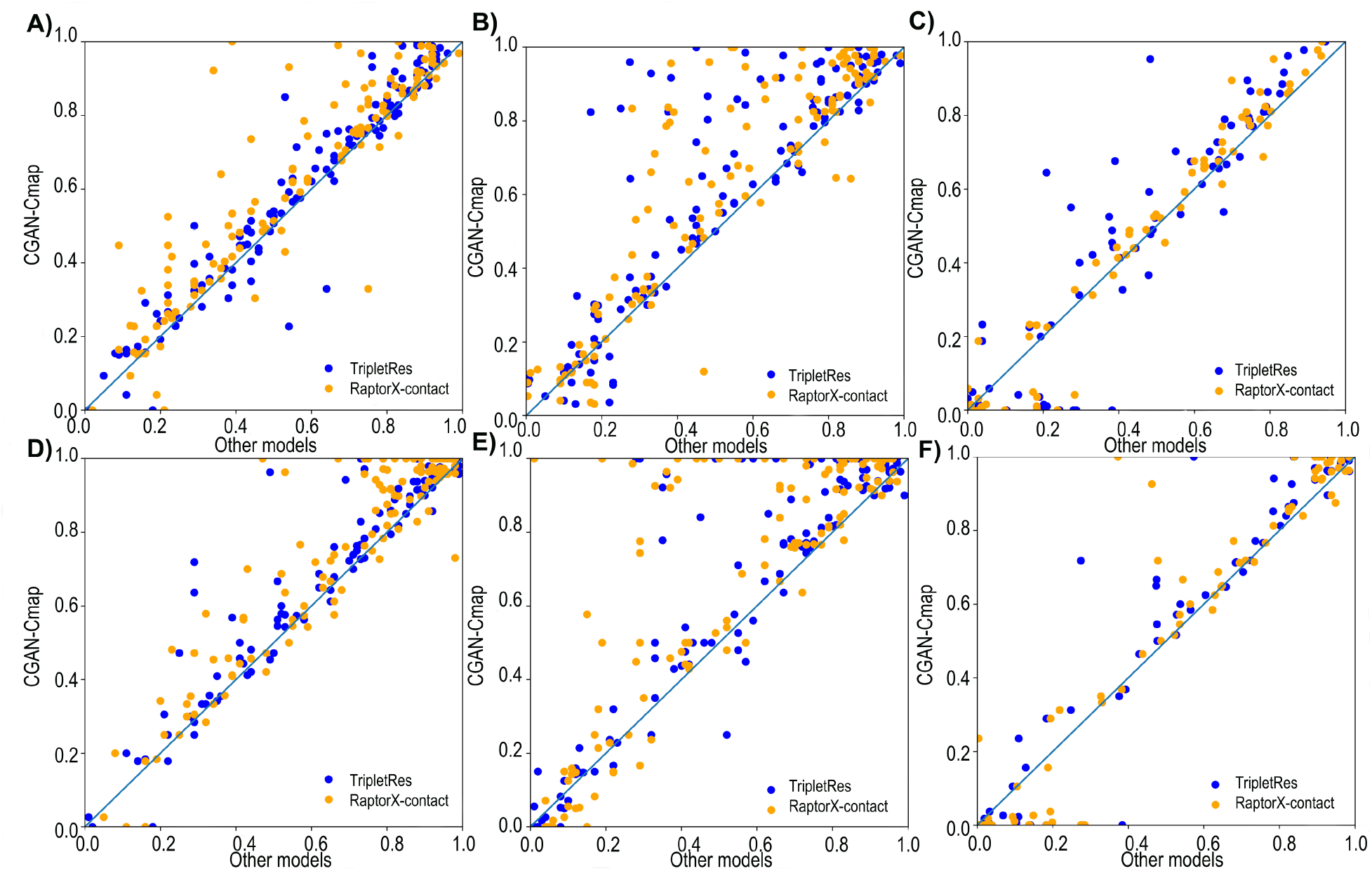
Performance comparison between CGAN-Cmap, TripletRes^17^ and RaptorX-contact^20^ for all individual targets in the CASP 12, CASP 13, and CASP 14 datasets, based on precision of A) top L/2 medium + long-range predicted contacts, B) top L/2 long-range predicted contacts, C) top L/2 extra long-range predicted contacts, D) top L/5 medium + long-range predicted contacts, E) top L/5 long-range predicted contacts, F) top L/5 extra long-range predicted contacts.

**Table 2.** Model performance of CGAN-Cmap, TripletRes^17^, RaptorX-contact^20^, DeepCon^45^, and DeepCov^46^ based on the mean precision of short, medium, and long-range top L/10, L/5, L/2, and L contacts in predicted contact maps for CASP 13 and CASP 14 test sets. The highest accuracy in each category is highlighted in bold font.

| Test sets | Model | (Medium + Long)-range |  |  |  | Long-range |  |  |  | Extra long-range |  |  |  |
| --- | --- | --- | --- | --- | --- | --- | --- | --- | --- | --- | --- | --- | --- |
|  |  | L | L/2 | L/5 | L/10 | L | L/2 | L/5 | L/10 | L | L/2 | L/5 | L/10 |
| CASP 13 | DeepCon | 0.56 | 0.70 | 0.81 | 0.86 | 0.42 | 0.54 | 0.64 | 0.71 | 0.33 | 0.47 | 0.62 | 0.65 |
|  | DeepCov | 0.46 | 0.58 | 0.69 | 0.74 | 0.33 | 0.43 | 0.53 | 0.60 | 0.28 | 0.40 | 0.53 | 0.61 |
|  | RaptorX-contact | 0.62 | 0.72 | 0.81 | 0.85 | 0.55 | 0.65 | 0.77 | 0.81 | 0.38 | 0.55 | 0.69 | **0.73** |
|  | TripletRes | 0.62 | 0.72 | 0.81 | 0.86 | 0.54 | 0.67 | 0.76 | 0.80 | 0.39 | 0.54 | **0.70** | 0.72 |
|  | CGAN-Cmap (standard) | 0.60 | 0.69 | 0.80 | 0.84 | 0.52 | 0.64 | 0.76 | 0.83 | 0.36 | 0.51 | 0.66 | 0.70 |
|  | CGAN-Cmap (dynamic) | **0.63** | **0.73** | **0.81** | **0.88** | **0.62** | **0.71** | **0.81** | **0.84** | **0.40** | **0.56** | 0.68 | 0.72 |
| CASP 14 | DeepCon | 0.28 | 0.37 | 0.45 | 0.56 | 0.16 | 0.26 | 0.32 | 0.38 | 0.17 | 0.22 | 0.25 | 0.28 |
|  | DeepCov | 0.21 | 0.27 | 0.32 | 0.47 | 0.13 | 0.20 | 0.29 | 0.34 | 0.11 | 0.13 | 0.16 | 0.21 |
|  | RaptorX-contact | 0.32 | 0.42 | 0.57 | 0.64 | 0.24 | 0.32 | 0.40 | 0.45 | 0.23 | 0.28 | 0.31 | 0.36 |
|  | TripletRes | 0.33 | 0.42 | 0.56 | 0.63 | 0.25 | 0.31 | 0.41 | 0.44 | 0.22 | 0.26 | 0.29 | 0.36 |
|  | CGAN-Cmap (standard) | 0.32 | 0.41 | 0.55 | 0.62 | 0.23 | 0.32 | 0.40 | 0.45 | 0.22 | 0.27 | 0.28 | 0.36 |
|  | CGAN-Cmap (dynamic) | **0.35** | **0.46** | **0.62** | **0.69** | **0.32** | **0.38** | **0.44** | **0.48** | **0.24** | **0.29** | **0.33** | **0.39** |

To demonstrate the applicability of our model, we present an example from the first domain of T1049 in CASP14 (PDB ID:6y4f) with 130 residues (Figure 3), a Zn-dependent receptor-binding domain of Proteus mirabilis MR/P fimbrial adhesin MrpH^47^. CGAN-Cmap has a mean precision of 88.2% for the top L/2 long-range contacts, compared to 57.8% by RaptorX-contact and 72.4% by TripletRes, respectively. As shown in Figure 3A and 3B, RaptorX-Contact and TripletRes do not predict any long-range contacts in Region 1, a critical loop-loop contact region. RaptorX-contact fails to predict contacts in Region 2 (Figure 3A). By contrast, CGAN-Cmap predicts both Regions 1 and 2, with high accuracy (Figure 3C).

**Figure 3.**
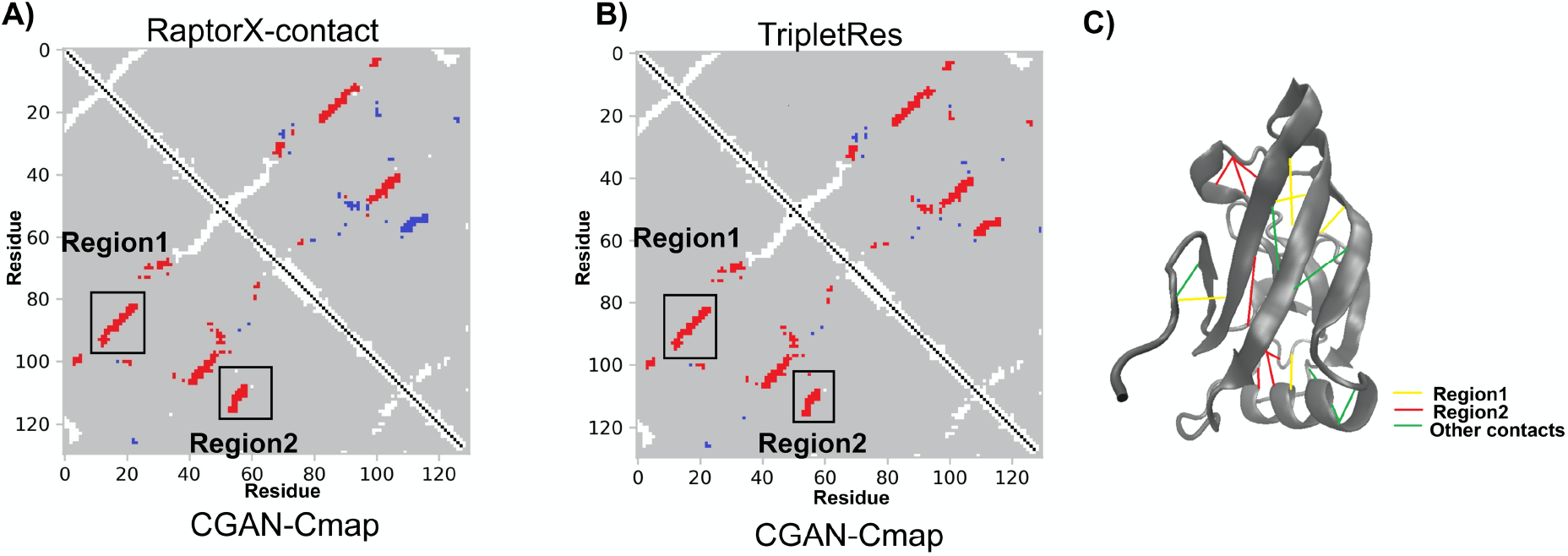
An illustrative example of a CASP14 target T1049 showing a comparison of top-L/2 long-range contact prediction by A) CGAN-Cmap (lower left triangle) and RaptorX-contact^20^ (upper right triangle) and B) CGAN-Cmap (lower left triangle) and TripletRes^17^ (upper right triangle). In each map, the true contacts are marked in white, true positives in red, and false positives in blue. C) Experimental structure of target T1049, with the long-range true positive prediction by CGAN-Cmap in Region 1, Region 2 and other regions marked with yellow, green, and red lines, respectively.

The performance of CGAN-Cmap is superior to existing state-of-the-art tools on CASP datasets, exceeding the best current methods by a large margin. This improved performance of CGAN-Cmap compared to other existing models originates from its dynamic weighted BCE loss function and its novel architecture. To summarize the effect of the architecture on the performance of the model, we highlight various blocks within the generator that are deterministic components of CGAN-Cmap driving accurate prediction of the contact map. First, our model uses a conversion block to convert protein 1D information to useful 2D feature maps for contact map generation. This block avoids repetition of information, and allows the model to learn the best converted 2D features from the input 1D information. Second, the synthesis blocks can effectively determine the essential parts of the protein 2D information that are useful for generating the contact map^48^. Finally, SE-Concat captures more information from the input feature maps because it not only reduces gradient vanishing owing to feature reusability but also highlights the most important information from feature maps, which results in the capture of complex sequence-contact relationships while using many fewer parameters than other methods^41^. All these blocks work together to generate state-of-the-art performance for contact map prediction. In addition to the unique combination of architecture blocks used in our model, our custom loss function overcomes the imbalance problem by effectively assigning the appropriate weights in the training process to predict highly sparse contacts. In the following sections, we discuss the effect of SE-Concat blocks and input features on the performance of CGAN-Cmap in detail, and we characterize the dependency of our model’s performance on the MSA generation protocol and number of homologous sequences within MSAs.

### SE-Concat block improves CGAN-Cmap performance to predict sparse contacts

One of the key novelties of our model is use of the SE-Concat block instead of the standard ResNet or SE-ResNet layer. To show the effect of SE-Concat on the performance of CGAN-Cmap, we calculate the performance of the model with SE-Concat and SE-ResNet, respectively, on CASP 13 and CASP 14 and summarize the performance in Table 3. Our CGAN-Cmap with SE-Concat blocks has at least 4%, and 6% mean precision improvement, for CASP 13 and CASP 14, respectively, on various top L/k long-range contacts compared to CGAN-Cmap with SE-ResNet blocks. We also calculate the average precision on the training set over the training epoch (Figure 4). These results confirm that CGAN-Cmap with SE-Concat displays a consistently better performance than CGAN-Cmap with SE-ResNet in most epochs of the training process. SE-Concat blocks help to extract more meaningful patterns from the input features, since the concatenation layer reuses the extracted features and helps to increase the information flow between layers^40,41^. The enhancement in information flow alleviates the vanishing-gradient problem and strengthens feature propagation to more accurately predict sparse contacts within the contact maps.

**Figure 4.**
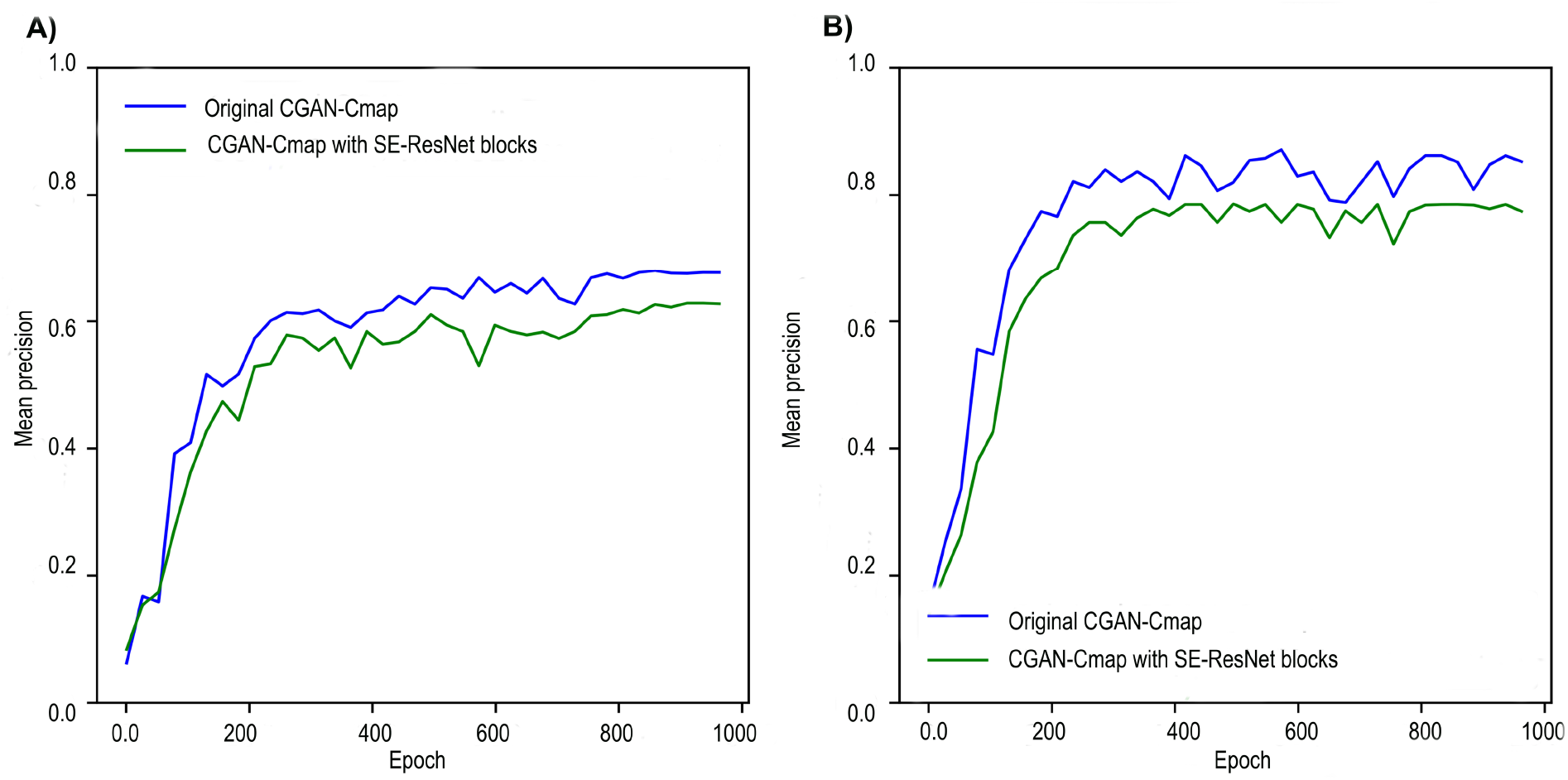
Comparison of the performance of the original CGAN-Cmap and CGAN-Cmap with SE-ResNet blocks by evaluation of the mean precision of the validation set over training epochs A) for top L/2 long-range contacts, and B) top L/5 long-range contacts.

**Table 3.**
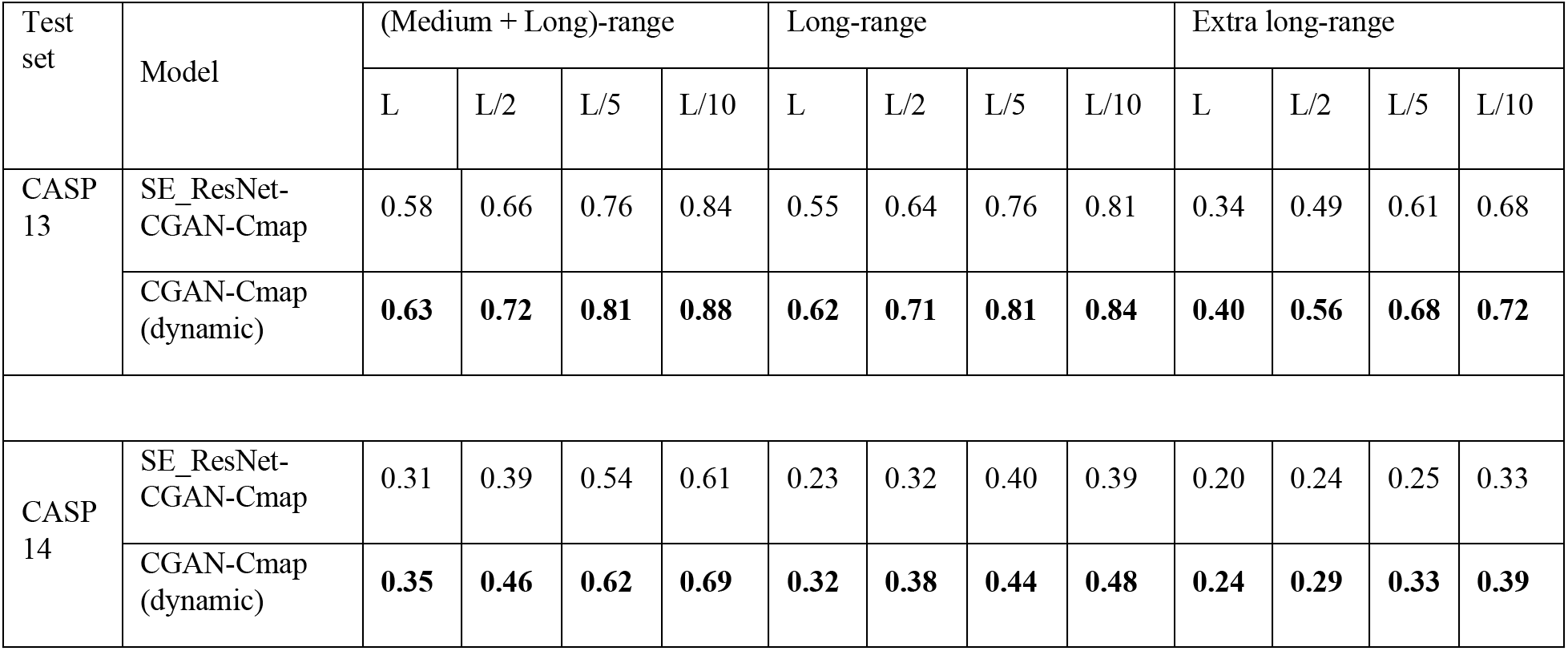
Model performance of the original CGAN-Cmap and CGAN-Cmap with SE-ResNet blocks instead of SE-Concat based on the mean precision of the predicted contact map on CASP 13 and CASP 14 test sets.

### Co-evolutionary pairwise features are key determinants in contact map prediction

Input features are one of the key determinants of model performance. To clarify the impact of individual groups of input features on the performance of the CGAN-Cmap model, we examine the efficiency and importance of the ensembled feature collection on the contact predictions. To do so, we categorize all input features into four groups: 1) sequential features (L x 54), 2) only pairwise features (L x L x 5), 3) pairwise features (L x L x 5) + SS3 & SS8 (L x 11) + ACC (L x 3), and 4) pairwise features (L x L x 5) + PSSM (L x 20) + PSFM (L x 20). In all groups, except the first, we consider pairwise features since we expect these to be essential input features for protein contact map prediction. Figure 5 presents the average top L/5 long-range contacts precision values over the training epochs. All models become stable after 300 epochs of training and reach a precision value of 65.4%, 71.9%, 80.4%, 82.3%, and 86.1%, for groups 1, 2, 3, 4, and an ensemble of all input features, respectively. CGAN-Cmap trained with an ensemble of all input features achieves consistently better performance than CGAN-Cmap trained with one of groups 1 to 4 only. In contrast to the model which uses sequential features (group 1), CGAN-Cmap with pairwise features (group 2) achieves higher mean precision for protein contact prediction, confirming that pairwise features are key elements for input. We note that when using group 4, our model achieves a mean precision close to the original model with an ensemble of all input features (lower by only 4%), suggesting that features derived directly from MSA such as PSSM and PSFM are critically important co-evolution features for improving performance of CGAN-Cmap. This analysis highlights the importance of pairwise features and specifically MSA-derived features (PSSM, PSFM) for contact map prediction, because these co-evolutionary features distinguish correlations resulting from direct or indirect effects of residue interactions, and thereby improve prediction accuracy of contacts. Result with group 3 suggest that other sequential features such as ACC and SS3 & SS8 can be used for further improvement of the model.

**Figure 5.**
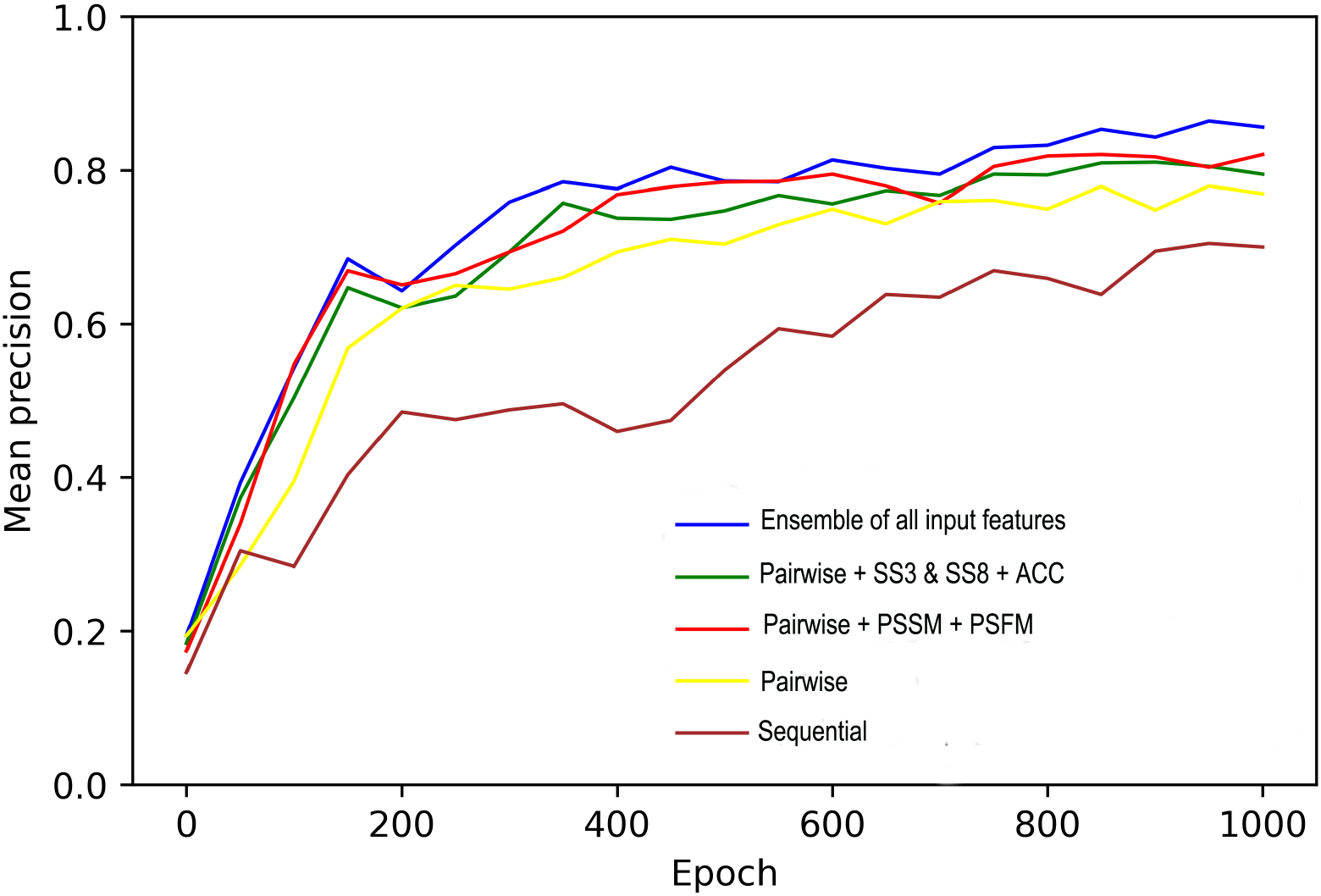
Mean precision of top L/5 long-range contacts over training epoch for the validation set showing the impact of various input features (ensemble of all input features, sequential features, pairwise, pairwise + PSSM + PSFM, and pairwise + SS3 + SS8 + ACC) on CGAN-Cmap performance.

### Effect of MSA protocol on CGAN-Cmap performance

The quality of the generated MSA has an essential impact on predicted contact maps because of its role in input feature generation. To show the type of dependency of CGAN-Cmap performance on the MSA extraction protocol and, more importantly, on the number of homologous sequences within the MSA, we train our model on input features extracted from MSA generated via only the HHblits approach^49^ without any constraint on Neff value and compare its performance with original CGAN-Cmap trained with DeepMSA (Table 4). The model’s performance with DeepMSA is only slightly better than with HHblits on different ranges of contacts (by at most 2.5%), suggesting that our model has a modest dependency on the MSA building protocol and even with HHblits or other MSA generation approach, the model still maintains its performance. To further validate this modest dependency, we compare the precision of top L/2 and top L/5 long-range contacts in predicted models with and without deep MSAs for all targets in CASP 12, 13, and 14 test sets (Figure 6).The performance of CGAN-Cmap with both MSA building approaches is similar for most targets within the CASP test sets. In another analysis, we characterize the dependency of our model on the Neff value. To that end, first we divide targets into low Neff targets (Neff < 10) and high Neff targets (Neff > 10) and then define a precision threshold of 50%. Then, we calculate the precision of Top L/5 long-range contacts for all individual targets within the CASP 12, 13, and 14 by CGAN-Cmap, TripletRes^17^, and Raptorx-Contact^20^(Figure 7). Our model predicts most of the high Neff targets with a precision greater than 50% (Figure 7A), confirming that higher Neff targets are predicted accurately with higher likelihood based on the higher quality information in their MSA. Interestingly, for low Neff targets, our model predicts about 67% (28/44) of them with the precision of more than 50%, suggesting that CGAN-Cmap can predict protein contact maps with high accuracy for proteins with a low Neff value. For comparison, we perform the same analysis for RaptorX-contact and TripletRes models (Figures 7B and 7C). TripletRes has good performance on high Neff targets, but it cannot predict low Neff targets with high accuracy, in contrast to our model that demonstrates its low dependency on number of homologous sequence within the MSAs for contact map prediction.

**Figure 6.**
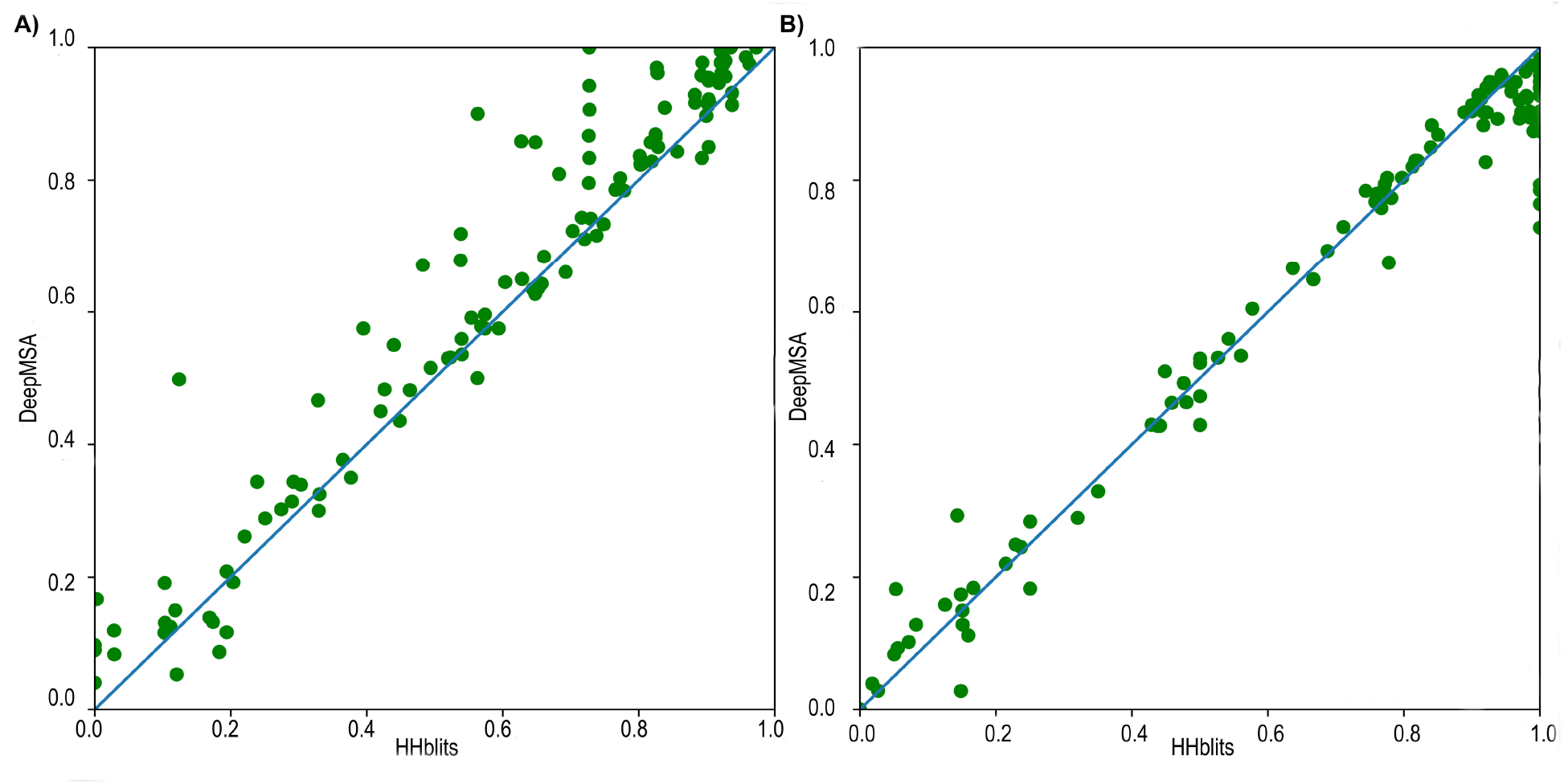
Precision of A) top L/2 long-range contacts and B) top L/5 long-range contacts predicted by CGAN-Cmap with deep MSAs versus that without deep MSAs for all targets in CASP 12, 13, and 14 test sets.

**Figure 7.**
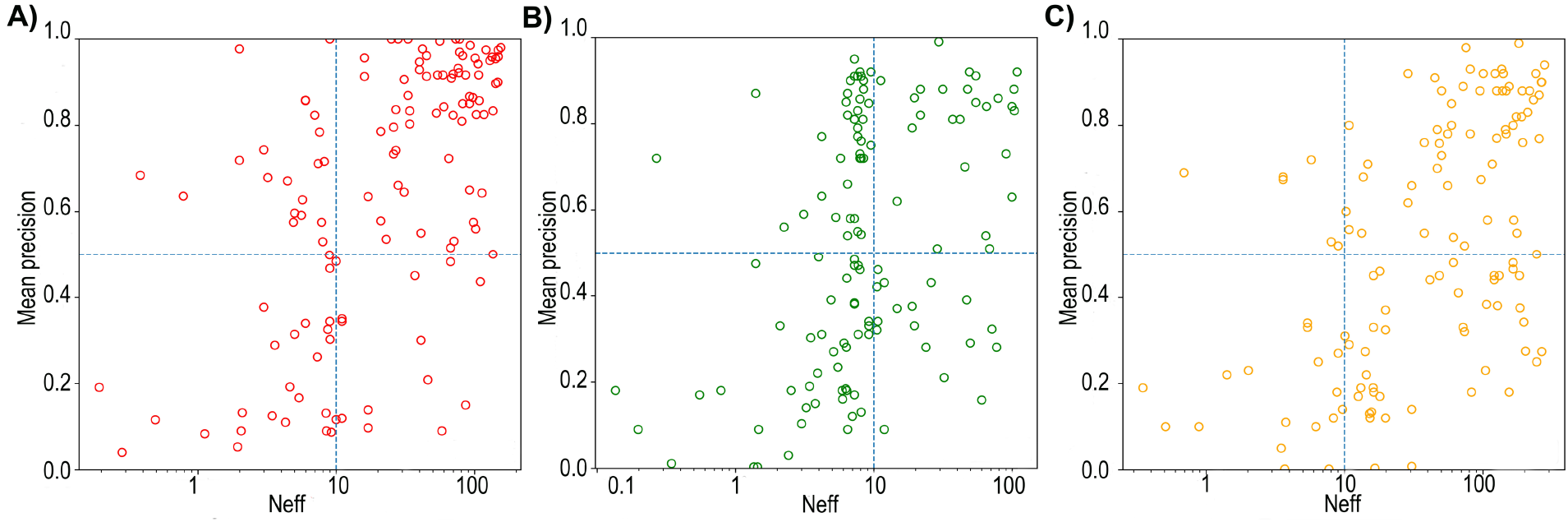
The precision of top L/5 long-range predicted contacts versus Neff of MSAs for A) CGAN-Cmap, B) RaptorX-contact^20^, and C) TripletRes^17^. Horizonal dotted line is precision threshold (50%) and vertical dotted line represents Neff threshold (10).

**Table 4.** Model performance of the original CGAN-Cmap with DeepMSA and CGAN-Cmap with HHblits based on the mean precision of predicted contact map on CASP 13 and CASP 14 test sets. The highest accuracy in each category is highlighted in bold font.

| Test set | Model | (Medium + Long)-range |  |  |  | Long-range |  |  |  | Extra long-range |  |  |  |
| --- | --- | --- | --- | --- | --- | --- | --- | --- | --- | --- | --- | --- | --- |
|  |  | L | L/2 | L/5 | L/10 | L | L/2 | L/5 | L/10 | L | L/2 | L/5 | L/10 |
| CASP 13 | HHblits-CGAN-Cmap | 0.62 | 0.70 | 0.80 | 0.86 | 0.60 | 0.68 | 0.79 | 0.83 | 0.42 | 0.53 | 0.66 | 0.69 |
|  | CGAN-Cmap (dynamic) | **0.63** | **0.72** | **0.81** | **0.88** | **0.62** | **0.71** | **0.81** | **0.84** | **0.40** | **0.56** | **0.68** | **0.72** |
| CASP 14 | HHblits-CGAN-Cmap | 0.33 | 0.44 | 0.59 | 0.66 | 0.30 | 0.37 | 0.42 | 0.47 | 0.23 | 0.26 | 0.32 | 0.37 |
|  | CGAN-Cmap (dynamic) | **0.35** | **0.46** | **0.62** | **0.69** | **0.32** | **0.38** | **0.44** | **0.48** | **0.24** | **0.29** | **0.33** | **0.39** |

## Conclusions

Accurate prediction of protein contact maps has been widely used in de novo protein structure prediction. In this study, we propose a novel GAN-based architecture, CGAN-Cmap, for contact map prediction. During the adversarial learning process, the generator network of CGAN-Cmap captures the underlying contact information from versatile protein features by employing a dedicated generator including various blocks such SE-Concat and synthesis blocks, while the discriminator network learns the difference between generated contact maps and real ones and automatically transfers them back to the generator network. By taking advantage of the novel architecture and training approach, our model outperforms all existing protein contact map predictors when the evaluation is assessed on different CASP targets. Furthermore, our proposed model exhibits a modest dependency on the number of homologous sequences in the MSA, resulting in accurate predictions for proteins with low Neff value. Although CGAN-Cmap shows promising performance for protein contact map prediction, there is still room for further improvement. In future studies advanced GAN training methods, such as WGAN^50^, can improve training stability. A symmetric loss functions such as the focal loss^51^ may also boost the predictive power of GAN-based architectures. Overall, the presented GAN-based deep learning architecture for contact map prediction can efficiently improve overall prediction performance and serves as a powerful new tool for contact map prediction.

