## Supplementary Information for "CGAN-Cmap: protein contact map prediction using deep generative adversarial neural networks"

### Dataset construction

We prepare our datasets with the following criteria:

1) sequence length is between 25-700 residues.

2) the maximum sequence identity is set to 30% to remove redundant sequences.

3) the native contact map resolution is more than 2.5 Å.

4) the protein contains a single chain.

Table S1. The number of proteins retained in each dataset construction step.

| Construction step | Training set | CAMEO | CASP 11 | CASP 12 | CASP 13 | CASP 14 |
|---|---|---|---|---|---|---|
| Input data | 11,593 | 76 | 127 | 72 | 41 | 51 |
| After criteria 1 | 10,648 | 76 | 110 | 72 | 41 | 49 |
| After criteria 2 | 8,969 | 76 | 110 | 72 | 41 | 41 |
| After criteria 3 | 7,902 | 76 | 110 | 59 | 41 | 40 |
| After criteria 4 | 7,274 | 76 | 110 | 54 | 41 | 40 |
| Overlap removal with test sets 25% identity | 7,128 | 76 | 110 | 54 | 41 | 40 |

**DeepMSA: MSA building approach**

To build a machine learning model, we need to extract some features from our input sequences. We extract both sequential and pairwise (co-evolution) features, in approach similar to that introduced in Raptorx-Contact[1]. The most important features are pairwise features captured from MSAs. To generate MSA for training and test sets, we use the modified version of DeepMSA first proposed by Li et al[2]. For the training set, we employ HHblits2[3] with three iterations, an E-value threshold of 0.001 and a minimum sequence coverage of 30% to search through the Uniclust30 (2017_10)[4] database to generate MSAs. First, we define Neff (Eq S1), representing the number of effective sequences in MSA, where $N$ is the total number of sequences in the MSA and $II\left[S_{m,n} > 0.8\right]$ is the conditional equation which is equal to 1 if the sequence identity $S_{m,n}$ between sequences m and n is greater than 0.8; or = 0 otherwise. We set the threshold of Neff to 80. In our initial MSA generation, we utilize the HHblits procedure used for the training set. If the Neff value is less than 80, we employ a second method for MSA generation, via jackhmmer[5] through searching UniRef90 (release-2017_12)[6]. If the Neff value still does not satisfy the given threshold, we employ a third MSA generation method by using hmmsearch[5] through the MetaClust (2017_05)[7]. For consistency of MSAs generated via different approaches, we convert all of them to HHblits format, then concatenate them. Figure S1 represent the MSA generation procedure.

$$Neff = \frac{1}{L}\sum_{n=1}^{N} \frac{1}{1+\sum_{m=1}^{N} II[S_{m,n}>0.8]} \quad \text{(Eq. S1)}$$

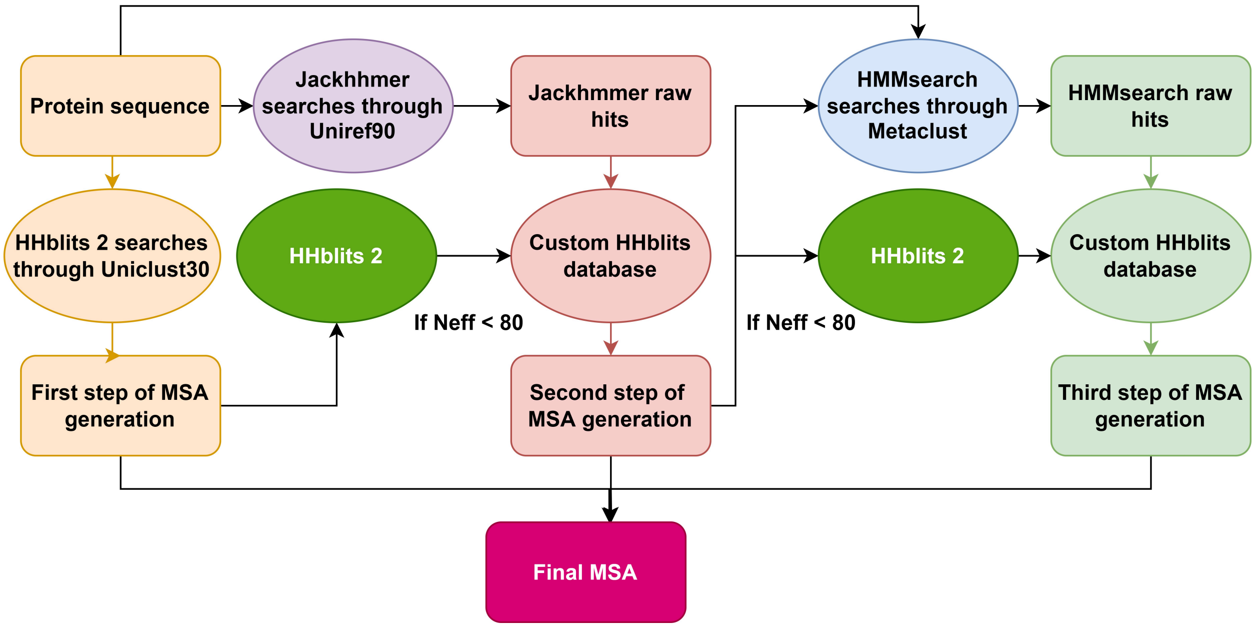


Figure S1. Modified DeepMSA method for MSA building from protein sequence.

### Model architecture and blocks

#### Custom blocks used within the model architecture

The generator of CGAN-Cmap consists of three subnets. The synthesis subnet includes a synthesis block (Figure S3A) that combines the feature maps of the 2D information with the previous layers and determines the information that should be kept for the next layers. The synthesis block separates half of the features extracted from the 1D information and the noise, then by using the sigmoid function, it can easily determine the more important parts of the separated features. By multiplying this information with the feature maps of the 2D information, the synthesis block determines which parts of the feature maps should be included in the contact map generation. The GANTL model[8] confirms that the synthesis block can boost the quality of the generated images.

The feature extraction subnet uses a series of SE-Concat blocks (Figure S3B) that help to extract the useful patterns from the input. The SE-Concat blocks are inspired by the SE-Res blocks introduced and used in DSResSol[9,10]. However, instead of the summation layer in that block, there is a concatenation layer that reuses the extracted features and helps to increase the information flow between layers[11]. The upsampling block (Figure S3C) is utilized to increase the current feature to something larger than the feature output of the next block for the model to learn more meaningful features within feature maps. The SE-Concat block contains the squeeze and excitation (SE) block (Figure S3D), which highlights the important information and improves the channel interdependencies[10]. Unlike the typical models which treat the feature maps equally, SE weighs the important information based on its effect on the final prediction, boosting the model performance[10].

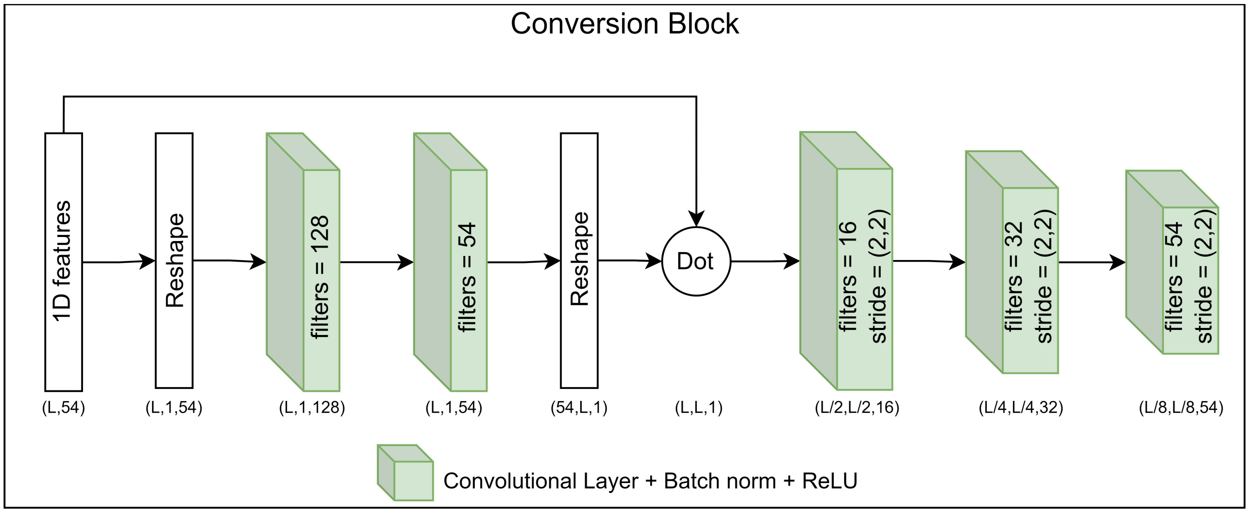


Figure S2. Conversion subnet architecture responsible for converting sequential (1D) features to 2D features.

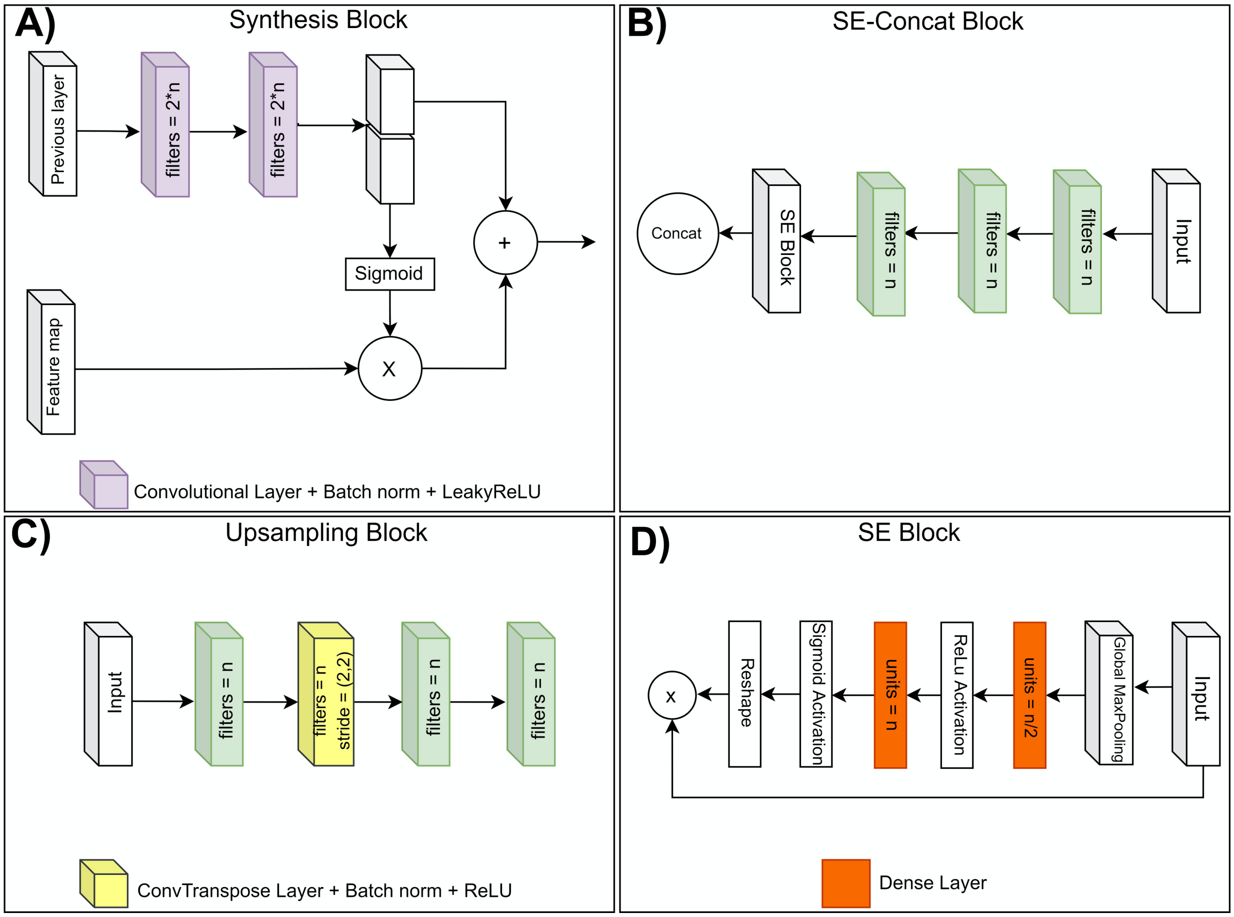


Figure S3. Architecture of 4 blocks used in the generator module: (A) synthesis block, (B) SE-Concat block, (C) upsampling block, and (D) SE block.

### Complete model architecture

Figure S4 shows the complete model architecture of CGAN-Cmap with size, number of units, and filters of each block.

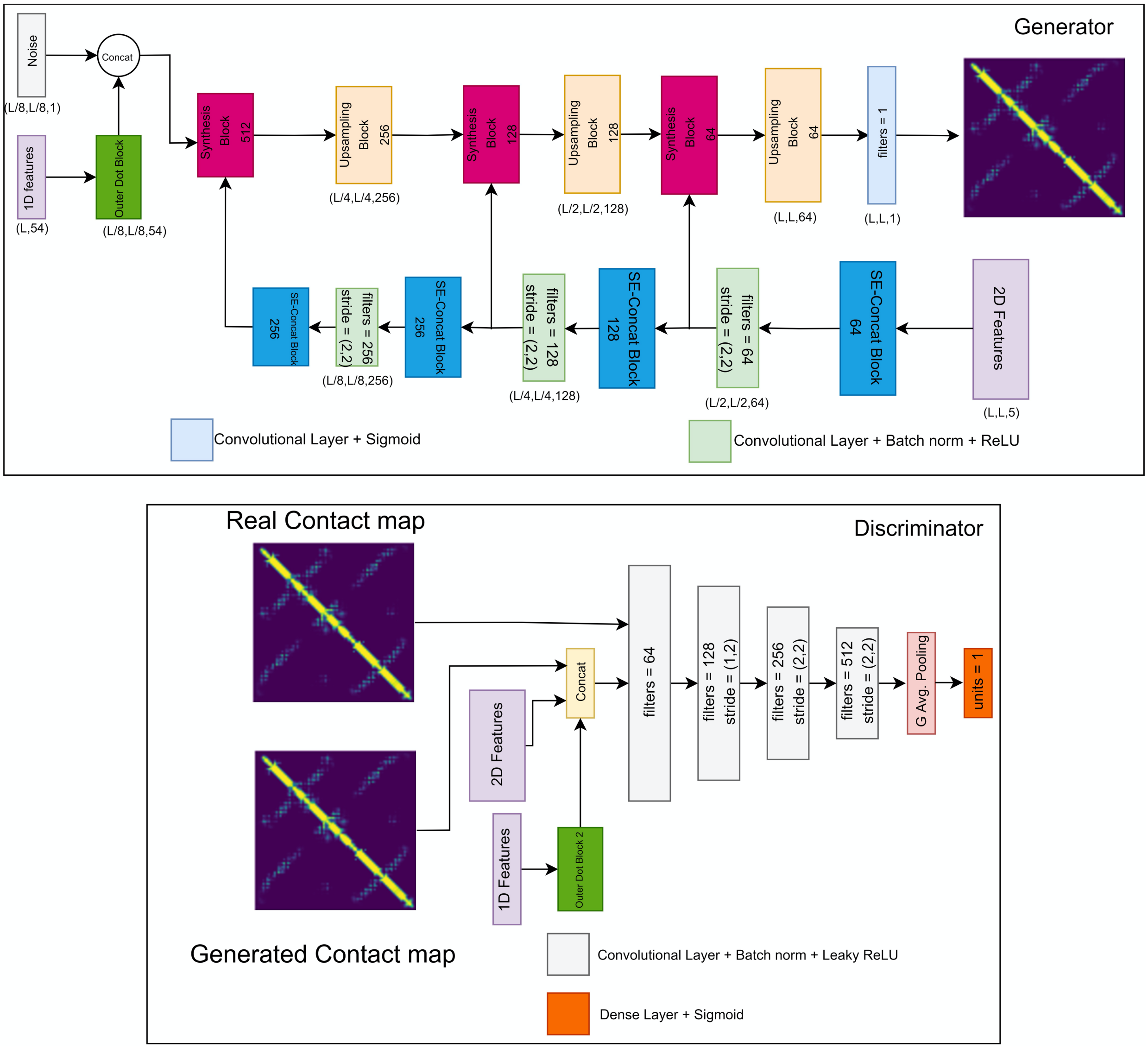


Figure S4. Architecture of CGAN-Cmap including architecture and size of each block within the generator and discriminator.

Table S2. Tested hyperparameters in network layers and optimal values generated via the Grid Search Method.

| Layers | Number of tested units or filters | Optimal Value | Filter size | Parameter s | Tested Values | Optimal Value |
|---|---|---|---|---|---|---|
| Number of SE-block | (6, 8, 10, 15) | 10 | - | Epochs | 1000 | 50 |
| Number of Synthesis block | (5, 8, 12) | 8 | - | Learning rate | (0.005,0.008,0.01, 0.02) | 0.008 |
| Number of filter size within blocks | (32,64,128,256, 512) | 32 | - | Batch size | (4,8,16) | 4 |
| Size of upsampling blocks | (32,64,128,256) | 32 | 1 | Decay rate | (10e -7,10e-8, 10e-9) | 10e-7 |
| Number of CNNs | (4,6,8,12) | 8 | {64, 128, 256, 512} | Early stopping value | (3,4,5,6) | 5 |

**Performance of the model on CASP 11, 12, 13, and 14**

We compare the range of precision of the predicted contact maps for CGAN-Cmap, TripletRes, and RaptorX-contact on targets in CASP 11, 12, 13 and 14 (Figure S4 A-D). The mean precision of CGAN-Cmap for all ranges of contacts are higher than TripletRes[12] and Raptorx-Contact[1] which further validates the promising performance of CGAN-Cmap on CASP 11, CASP 12, CASP 13, and CASP 14 targets.

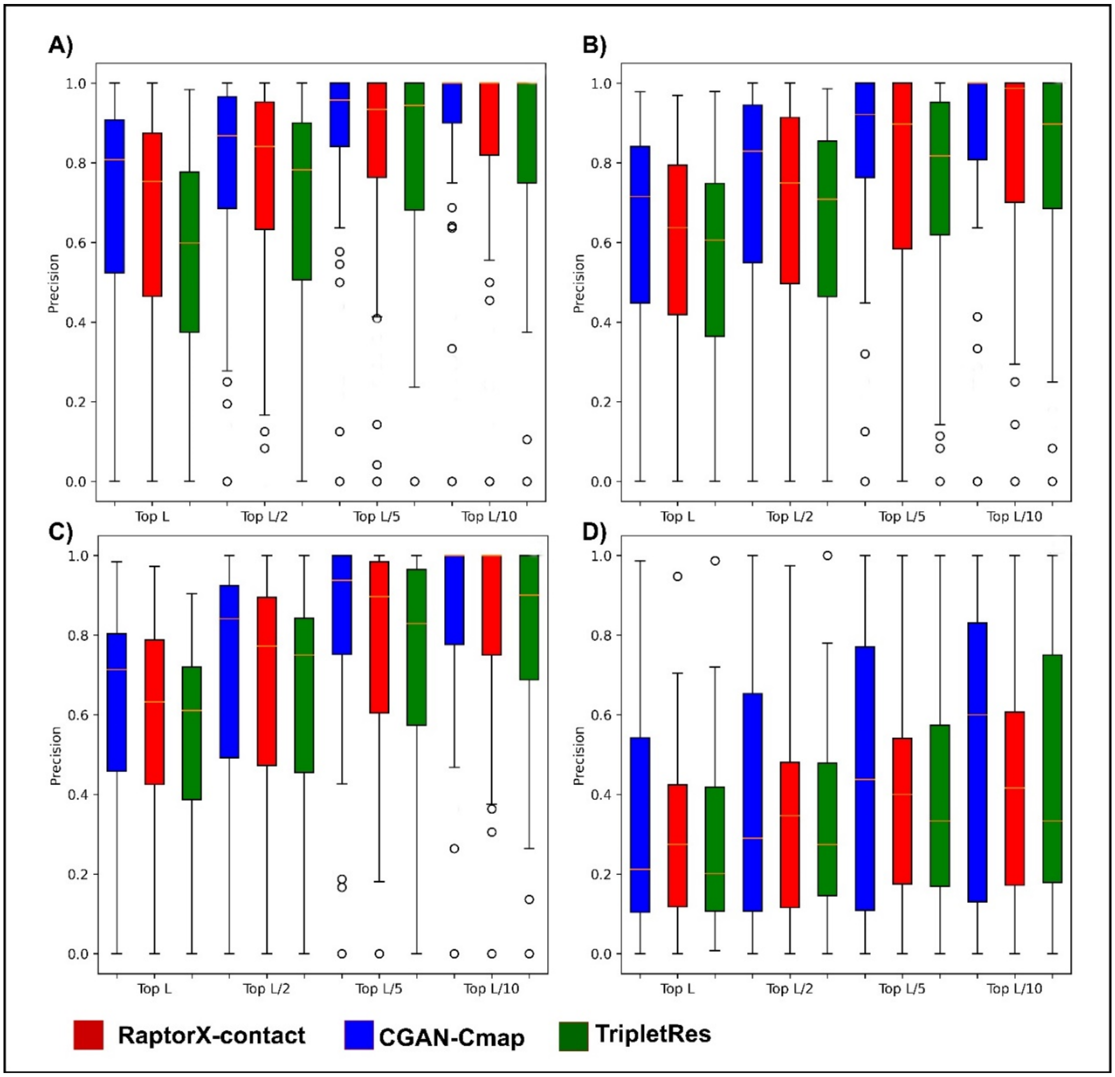


Figure S4. Box plots for performance of CGAN-Cmap, TripletRes[12], and Raptorx-Contact[1] on CASP test sets: A) CASP 11, B) CASP 12, C) CASP 13, D) CASP 14.

10 Hu, J., Shen, L. & Sun, G. in *Proceedings of the IEEE conference on computer vision and pattern recognition.* 7132-7141.

11 Huang, G., Liu, Z., Van Der Maaten, L. & Weinberger, K. Q. in *Proceedings of the IEEE conference on computer vision and pattern recognition.* 4700-4708.

12 Li, Y., Zhang, C., Bell, E.W., Zheng, W., Zhou, X., Yu, D.J. and Zhang, Y. Deducing high-accuracy protein contact-maps from a triplet of coevolutionary matrices through deep residual convolutional networks. *PLoS computational biology* **17**, e1008865 (2021).
